# Adversity magnifies the importance of social information in decision-making

**DOI:** 10.1101/061630

**Authors:** Alfonso Perez-Escudero, Gonzalo G de Polavieja

## Abstract

In adverse conditions, individuals follow the majority more strongly. This phenomenon is very general across social species, but explanations have been particular to the species and context, including antipredatory responses, deflection of responsibility, or increase in uncertainty. Here we show that the impact of social information in realistic decision-making typically increases with adversity, giving more weight to the choices of the majority. The conditions for this social magnification are very natural, but were absent in previous decision-making models due to extra assumptions that simplified mathematical analysis, like very low levels of stochasticity or the assumption that when one option is good the other one must be bad. We show that decision-making in collectives can quantitatively explain the different impact of social influence with different levels of adversity for different species and contexts, including life-threatening situations in fish and simple experiments in humans.

---

Decision-making theory explains our tendency to follow the majority: when the choices of others inform about the quality of the options, the majority option has the highest estimated quality [1–4]. Less intuitive is that the probability to follow the majority should increase in adverse conditions. Yet this is exactly what happens to many social animals: when conditions deteriorate individuals follow each other more strongly, choosing the majority option with higher probability (Figure 1a). This phenomenon— which we will call *superaggregation in adversity*—has been extensively studied as a response to predators [5–10], but may also happen in other adverse situations, such as in the absence of food [11]. In humans, the occurrence of sudden bank runs [12–14] and human stampedes [15–21] suggests superaggregation in adversity, although data are insufficient to draw definitive conclusions. Different theories explain superaggregation in adversity in particular contexts, such as when aggregation provides protection against predators [10, 22–25], when adversity correlates with uncertainty [26, 27] or, in humans, when copying others’ behavior can help to deflect the responsibility for an anticipated failure [28, 29]. In contrast, we hypothesise that superaggregation in adversity is rooted in the basic structure of decision-making, being a general phenomenon across species and contexts.

Most studies dealing with superaggregation in adversity look for the optimal level of aggregation given a particular situation. For example, whatis the optimal level of aggregation when a predator attacks the group? By doing so, they implicitly neglect the limitations of individual decision-makers, whose actions are constrained by the information available to them. For example, a given individual may not be certain whether a predator is approaching the group or not. This individual-centered approach is the point of view of decision-making theories [1–4, 27, 30–34], but it has never been applied to superaggregation in adversity. We find that decision-making plays a key role, naturally leading to superaggregation in adversity in a wide range of situations.

Let us consider for example a person choosing between two identical doors (*x* and *y*) to find the exit from a building (Figure 1b). The deciding individual may have private information about the two doors (e.g. from previous experience) and social information (the behavior of other people). These two sources of information must be combined to assess the situation and make the best possible decision.

We divide the decision in two steps. First, the deciding individual estimates the quality of the available options. We will use “quality” as a generic name for whatever is relevant to the decision. In the context of evolution, quality will measure the fitness value of each option (for example, the quality of a particular escape route from a predator may be the probability to survive). In the language of economic decision-making, quality refers to the utility of each option. In the two-door example, we may define quality as the inverse of the time needed to reach the street. The qualities (*Q*_*x*_ and *Q*_*y*_) are estimated by combining the privately estimated qualities (*G*_*x*_, *G*_*y*_) with social information. We will consider symmetric private information, so the privately estimated quality is the same for both options (*G*_*x*_ = *G*_*y*_ = *G*). In the case of a group attacked by a predator, we can apply this analysis on several escape routes that seem equally good. In our example with two doors, private information will be symmetric if the decision-maker is unfamiliar with the building and both doors look identical. For two options, we can visualize the estimated qualities in a 2-dimensional quality landscape (Figure 1c, left). In the second step, a decision rule transforms the estimated qualities into the probability of choosing each option (Figure 1c, right).

Now let’s investigate the effect of adversity on the decision-making process. We define adversity as a decrease in the apparent quality of all options. For example, the approach of a predator creates an adverse situation, since the probability that an animal dies after choosing any given direction is higher than when the predator was absent. In our two-door situation, a fire in the building would create an adverse situation. We assume that the deciding individual perceives (at least partially) this deterioration, so the privately estimated qualities decrease accordingly. Note that we do not need to assume that the situation becomes more uncertain in adversity: the privately estimated qualities may be as accurate as they were before—just lower.

Adversity thus moves the estimation to a different point of the quality landscape, even if the social information remains constant (Figure 1d). In general, this new location corresponds to a different probability of choosing each option (Figure 1e, top). The only exception is when the estimated qualities run along an isoprobability line of the decision rule (Figure 1e, middle). Such a perfect match is unlikely except for decision rules with large regions of constant probability, as for example the deterministic rule “always choose the option with highest estimated quality” (Figure 1e bottom). This decision rule is used in many simple models, preventing them to explain superaggregation in adversity [1, 2]. Also, many current models assume mutually excluding options (one of them is good and the other is bad, so *P*(*x* is good) = 1 − *P*(*y* is good)) [1–3]. These models lack a parameter that measures adversity, which would require lowering *P*(*x* is good) and *P*(*y* is good) simultaneously, and cannot predict superaggregation in adversity.

In general, decision rules are not deterministic and options are not mutually excluding, so the probability of following the majority will change with adversity. But why do we usually observe superaggregation in adversity rather than subaggregation? We have found three different mechanisms that promote superaggregation in adversity.

## (a) Relative decision rules

Many decisions depend on the relative value of estimated qualities, rather than on their absolute value. Examples range from bacteria to humans, including rules such as Weber’s law, probability matching, and others [3, 4, 35–44]. All relative rules have isoprobability lines identical to those in Figure 2a (Methods). With these rules, when the social information remains the same (Figure 2a, green arrows) its impact increases in adversity, with a higher probability of following the majority (Figure 2b). Even when the influence of social information in the qualities decreases in adversity, the probability to follow the majority will increase as long as the estimated qualities fulfil

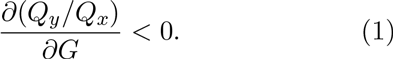

where *Q*_*x*_, *Q*_*y*_ are the final estimated qualities and *G* is the privately estimated quality (Methods). This condition is fulfilled both by non-optimal estimation rules (such as the one in Figure 2a) and by optimal ones, such as the Bayesian model shown in Figure 2c (Methods). This model—which reproduces experimental data of several species [4, 45, 46]—fulfils Equation 1 for any combination of parameters, predicting superaggregation in adversity in all conditions (solid line in Figure 2d; Methods). Other implementations of Bayesian decision-making produce the same result(section 1 of Supplement).

**Figure 1:**
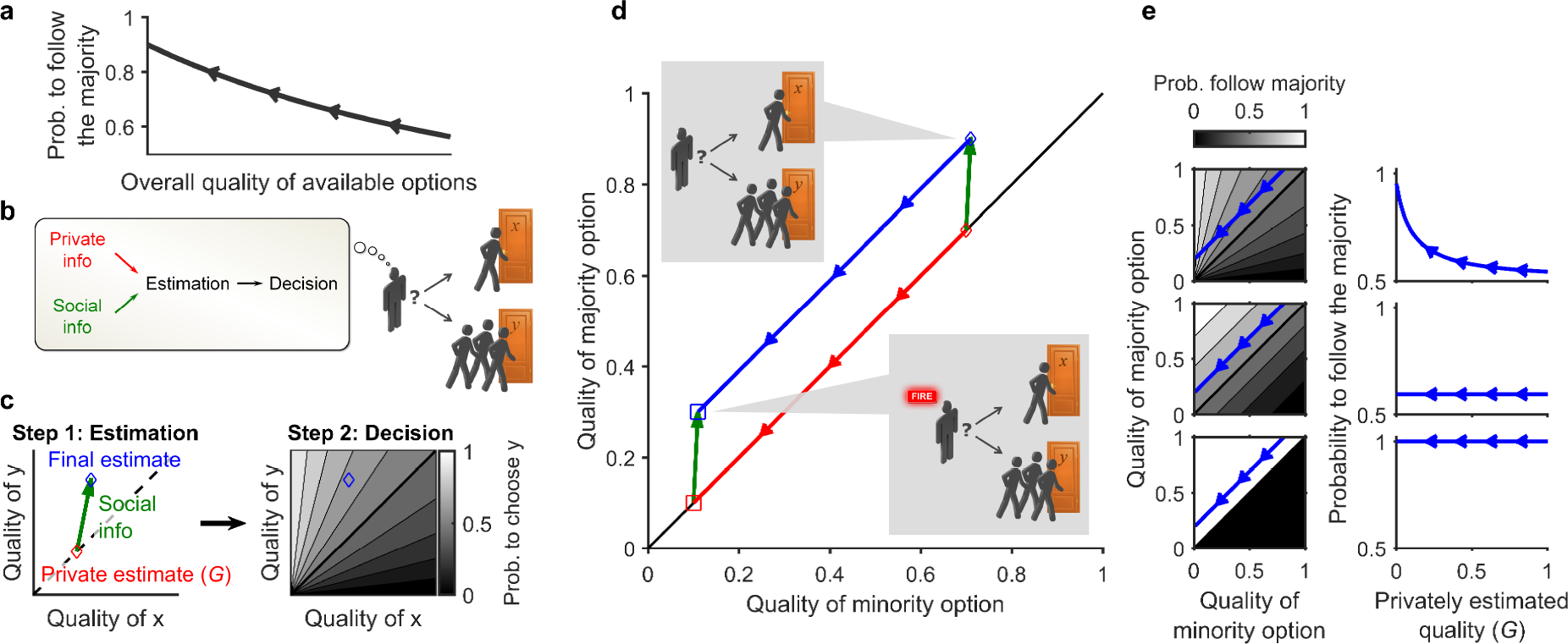
*Decision-making predicts that aggregation changes in adversity.* ***a**, Definition of superaggregation in adversity: the probability to choose the majority option increases as the quality of all options decreases. Arrowheads point in the direction of increasing adversity. **b**, A subject must choose between two seemingly identical options (doors x and y). One other person is choosing door x, and three other people are choosing door y, which is therefore the majority option. **c**, Decision process: In step 1, social and non-social information are integrated to estimate the qualities of the options, Q_x_ and Q_y_. Red diamond: privately estimated qualities (G). Green arrow: contribution of the social information. Blue diamond: Estimated qualities. In step 2, the decision rule gives the probability of choosing each option (P_x_ and P_y_) given the estimated qualities from step 1. In this example, P_y_ = Q_y_/(Q_x_ + Q_y_) and P_x_ = 1− P_y_. **d**, Example of the evolution of the estimated qualities when conditions become adverse. Colours as in c. Diamonds: favorable conditions. Squares: adverse conditions. Blue and red arrowheads point in the direction of increasing adversity. e, **Left**: Example of the trajectory followed by the estimated qualities in adversity (arrowheads point towards higher adversity), for three difierent decision rules: **top**: relative decision rule (P_y_ = Q_y_/(Q_x_ + Q_y_)); middle: absolute decision rule 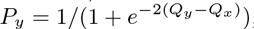 **bottom**: deterministic decision rule (P_y_ = 1 if Q_y_ > Q_x_, and P_y_ = 0 otherwise). **Right**: Probability to follow the majority as a function of the privately estimated quality, for the same trajectories and decision rules shown on the left.*

## (b) Saturation near the upper bound of qualities

The range of qualities is often bounded, for example when there is a limit in the amount of food an animal can consume or when qualities are probabilities. In this case, as the privately estimated qualities increase, both the private and the final estimates converge to the upper bound, so the difference between them—the contribution of social information—tends to zero. Thus, the same amount of social information can have a greater impact in adversity, when the estimation is further away from the upper bound (Figure 2c). To decouple the effects of saturation and relative decision rules, we considered a decision rule that depends on the difference of the qualities, *Q*_*y*_ − *Q*_*x*_ (Methods). With this decision rule, Bayesian estimation predicts superaggregation in adversity when the initial estimated qualities are high (dashed line in Figure 2d, see Methods). In the range of very low qualities we find the opposite effect, due to the lower bound of the probability. However, social information usually pushes the estimation towards the upper bound, making the range for superaggregation in adversity wider than the range for the opposite effect, especially for large groups (Methods).

## (c) Worsening of the worst-case scenario

Adversity can also modify the values of the potential payoffs behind each option, rather than the probability of reaching them. In our example of two doors in an unfamiliar building, a fire alarm does not change the probability that each door leads to the street, nor the corresponding payoff *r*_high_. Instead, it decreases the payoff of choosing the wrong door from a low value *r*_low_ (detour to street) to an even lower payoff 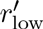 < *r*_low_ (possibility of dying in a fire). Adversity thus increases the contrast between a good and a bad choice, effectively rescaling the estimation from the region between *r*_low_ and *r*_high_ to a wider region of the quality landscape between 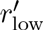 and *r*_high_ (Figure 2e). This rescaling results in a larger contribution of the social information, leading to superaggregation in adversity for both relative and absolute decision rules (Figure 2f) and for all combinations of the parameters (Methods). The opposite effect takes place when adversity affects the gains of a good option, rather than the cost of a bad one (Methods). Therefore, among adverse situations, we expect the life-threatening ones—characterized by a deterioration of the worst-case scenario—to produce stronger superaggregation in adversity.

**Figure 2:**
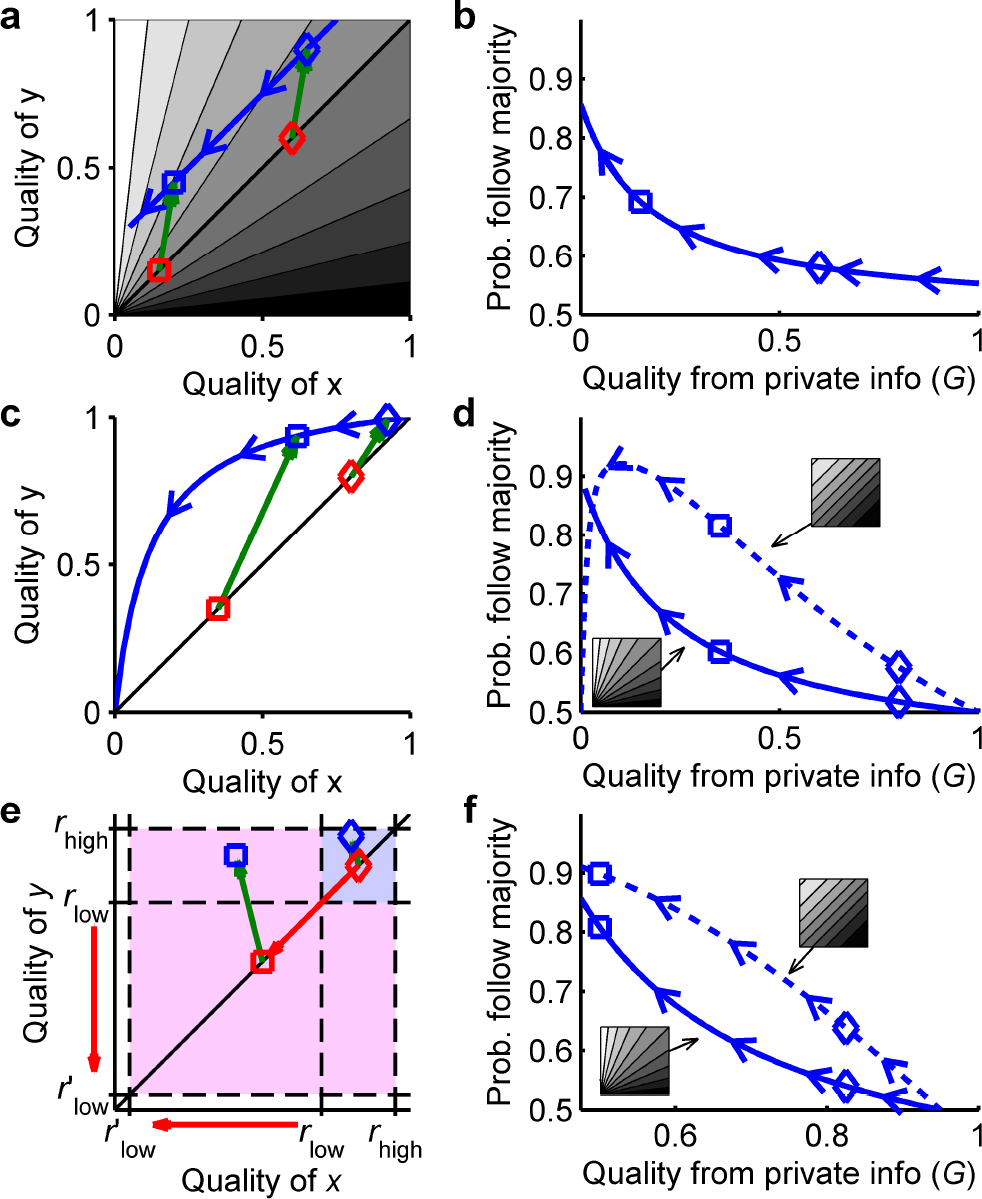
*Mechanisms responsible for superaggregation in adversity.* ***a**, Estimated qualities for an additive estimation model (Q_x_ = G + S_x_, Q_y_ = G + S_y_, with S_x_ = 0.05, S_y_ = 0.3). Red: private estimate for favourable conditions (G = 0.6, diamond) and adverse conditions (G = 0.15, square). Green arrows: contribution of the social information. Blue: trajectory of the final estimated qualities when adversity increases (arrows point towards adversity). Background: probability of choosing y for each pair of qualities, for the relative decision rule P_y_ = Q_y_/(Q_x_ + Q_y_) (grayscale: black=0, white=1). **b**, For the model in **a**, probability of choosing the majority option (y), as a function of the privately estimated quality (G). Arrows point in the direction of adversity. Diamond and square correspond to the same conditions as in a. c, Estimated qualities for a Bayesian decision-making model (Equation 17). Colours and symbols as in a. d, Probability of choosing the majority option (y) as a function of the privately estimated quality (G), using the Bayesian estimation model in c and a relative decision rule (P_y_ = Q_y_/(Q_x_ + Q_y_), solid line) or an absolute decision rule 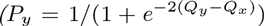 dashed line). e. Estimated qualities for the model with payoffs (Equation 22). Thigh is the payoff of a good option, r_low_ the payoff of a bad option in favourable conditions, and 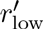 the payoff of a bad option in adverse conditions. Colours and symbols as in **a**. **f**, Same as **d** but for the model in e. Quality from private info is defined as G = r_low_ + (r_high_ − r_low_)P_private_, where P_private_ = 0.5 privately estimated probability that options have a high reward.*

**Figure 3:**
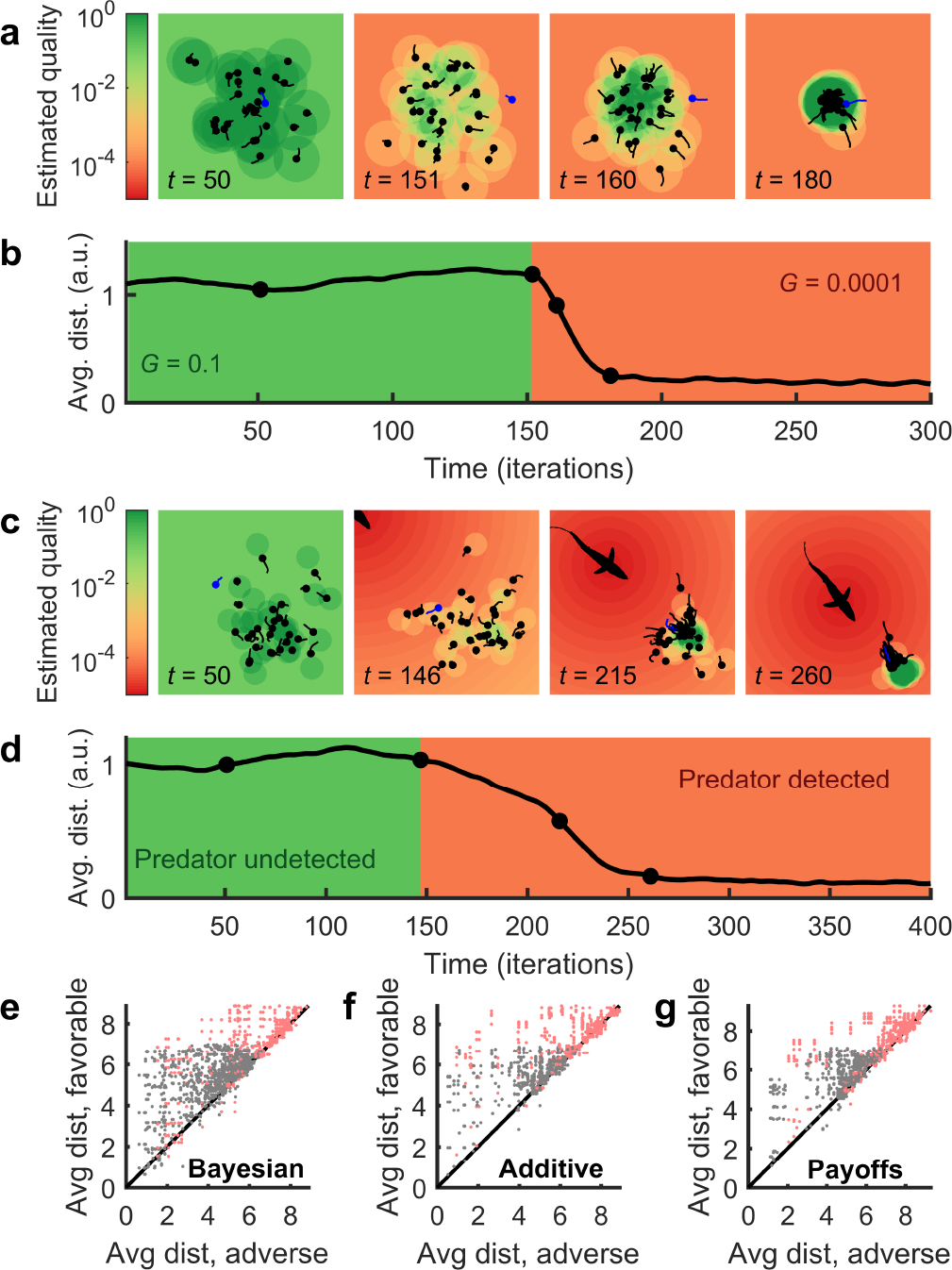
*Spatial superaggregation in adversity.* ***a**, Four frames of a simulation where private information is identical for all locations and changes suddenly at time t =150 from favourable conditions (G = 0.1) to adverse conditions (G = 0.0001). Each point is an individual. We chose an individual (blue) to illustrate its estimated probability that each pixel is a good location (background colour). **b**, Average distance among individuals as a function of time, for the same simulation as in a. Background colour indicates the value of G. Dots mark the times of the frames shown on **a**. **c**, Same as a, for a simulation with a predator. **d**, Same as **b**, for a simulation with a predator. **e**, Average distance between 10 individuals when conditions are favorable, vs. their average distance when conditions are adverse (points above the diagonal indicate superaggregation in adversity). Simulations use the same dynamical model as above, and a Bayesian decision rule. Each point comes from a random draw of all parameters of the model. Grey: simulations in 2D. Red: simulations in 3D. **f**, Same as e, but with an additive decision rule. **g**. Same as e, but with a rule with payoffs (Equation 22).*

Each of these three mechanisms can produce superaggregation in adversity by itself, but in general we may observe several of them reinforcing each other, as for example the solid lines in Figure 2d,f.

To illustrate superaggregation in adversity in a more general setting, we built a spatial model where individuals estimate the quality of all points within a given distance (see section 3 of Supplement for details). Social information increases the quality of the space around each individual. In favorable conditions, a point close to other individuals has only slightly higher quality than the rest of points, so the animals remain relatively disperse. When conditions deteriorate, estimated quality is high only for high density regions, so all individuals converge towards them (Figure 3a,b and Movies S1, S2). The model also shows aggregation when a predator approaches the group. Any individual that perceives the predator updates its private information to a non-uniform map of qualities whose minimum is at the predator’s position. This gradient pushes the individuals away from the predator and decreases the overall quality, increasing aggregation (Figure 3c,d and Movie S3). These results are independent of the mechanism behind superaggregation and of the model’s dynamical parameters: we have run simulations with different estimation models, in 2D and 3D, and with random parameters of the dynamical model (speed, acceleration, etc). Superaggregation in adversity arises often, independently of these details (Figure 3e,f,g).

We have also confronted our model with experimental data of groups of ten fish (*Fundulus diaphanus*) [11]. To modify the private information of the fish, the authors sprayed different odours uniformly on the water. Odours that signal adversity increased the cohesion in the fish (Figure 4a, insets). Our spatial model reproduces these data, keeping all parameters constant across conditions except private information *G* (Figure 4a).

Superaggregation in adversity may also take place in humans, as suggested by the occurrence of sudden bank runs [12–14] and human stampedes [15–21]. It is however unclear whether these events actually emerge from superaggregation, to other causes such as sudden changes in the private information towards a single preferred option, or to a combination of both. Tests in these panic situations are difficult, but our theory suggests that superaggregation in adversity should also happen when adversity is unrelated to life-threatening situations. In such situations our prediction is easy to test, and we did so using an existing dataset [47]. The experimental subject had to choose one out of eight face-down cards, and got a reward when the chosen card was red. The subject knew the proportion of red cards (so had private information about the probability of success), and could either choose one card or rely on the opinion of another person—either a professor or a student (Fig. 4b). The data show the trend predicted by our theory: Subjects rely more on social information the lower is the proportion of good cards (Figure 4c; Methods).

**Figure 4:**
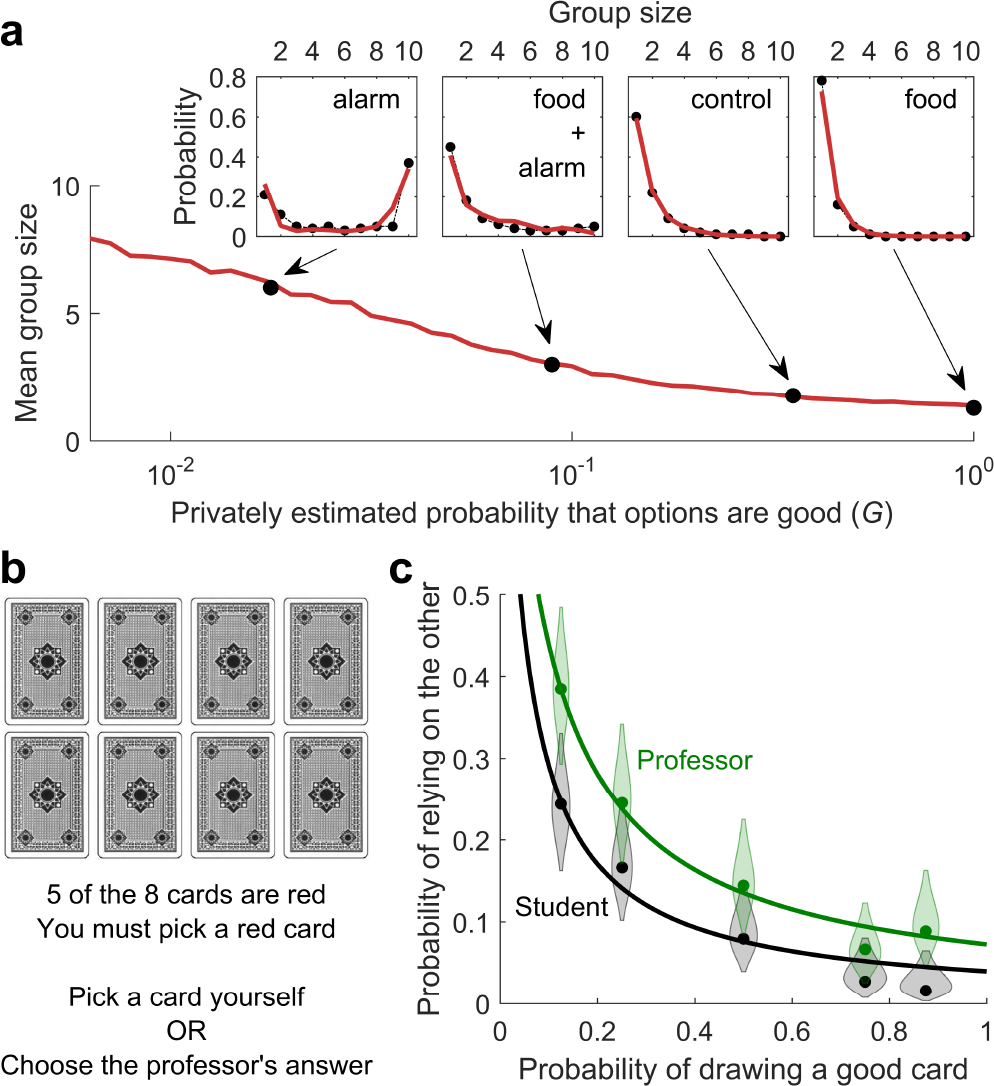
*Experiments are consistent with superagre-gation in adversity emerging from decision-making.* ***a**,Mean group size for 10 fish swimming in a closed space as a function of the privately estimated quality, as predicted by the spatial model. The black dots indicate the experimental mean group sizes [11], and the values of G that best fit the experiments. Insets: Distributions of group sizes for the four experiments (dots: experimental data. red line: simulations). **b**, Layout of the experiment of ref. [47]. **c**, Probability of relying in the card choice of another person (a professor for the green data and a student for the black data), as a function of the proportion of red cards (which is equivalent to the probability of getting a good card at random). Dots: experimental data (curved patches indicate experimental uncertainty due to sampling, with width proportional to the probability that the true value falls at each value, and truncated at the 95% confidence interval). Lines: model (Equation 28).*

## Discussion

We have shown that aggregation should change in adversity, and proposed three general mechanisms that make it increase rather than decrease. These mechanisms emerge from the basic structure of decisionmaking, explaining the generality of known cases of superaggregation in adversity [5–9, 11]. Furthermore, our results suggest that superaggregation in adversity may be a general feature of decisions in social contexts, and should be investigated in a wider range of situations.

Our results are complementary to other mechanisms that may increase aggregation in adversity in specific situations [22–25], and in fact our framework may incorporate many of these models. For example, it can reproduce the selfish-herd hypothesis[23] when the qualities are given by the probability of surviving a predator attack (see section 2 of Supplement and Figure S1).

All our results apply to the case when the presence of other individuals in one particular option increases its expected quality. This will not be the case if competition is strong, outweighing the effect of social information. In this case we might expect different— perhaps even opposite—trends. We therefore expect superaggregation in adversity to be strongest when competition is weak.

Our analysis has focused on symmetric private information (all options have the same privately estimated quality). The mechanisms discussed are also at work with asymmetric private information, and will typically increase aggregation around the privately preferred option (when it is the same for all individuals). In this case it is difficult to decouple the role of private and social information, but superaggregation will usually take place. An additional confounder arises if the asymmetry in private information between the options increases or decreases in adversity, obscuring the effect of social information. For example, many situations are characterized by a sharp increase in asymmetry: people wandering in a building usually have different aims, but a fire alarm will direct everyone towards the exits. This type of aggregation is compatible with our results and fits in our framework, but our results are applicable even when considering symmetric options, as for example two identical emergency exits. In general, by sticking to the symmetric case, we have shown that superaggregation in adversity does not require any asymmetry in the private information.

Superaggregation in adversity is at the core of decision-making. This finding does not exclude other causes for the observed behaviors, but should be taken as a baseline before resorting to less parsimonious explanations.

## Methods

### Formal definition of superaggregation in adversity

For a given decision-making model, we will say that superaggregation in adversity occurs whenever

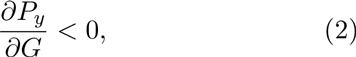

where *P*_*y*_ is the probability to choose the majority option, and *G* is the privately estimated quality (equal for all options because we consider symmetric private information).

### Relative decision rules

Relative decision rules are those that depend on the relative values of the estimated qualities, rather than on their absolute values. This family of decision rules includes those that depend on the ratio of two qualities, *Q*_*y*_/*Q*_*x*_ [35], on the relative difference between qualities, (*Q*_*y*_ − *Q*_*x*_)/(*Q*_*y*_ + *Q*_*x*_), such as Weber’s law [36–38], and on the relative value of qualities, *Q*_*y*_/(*Q*_*x*_ + *Q*_*y*_), such as probability matching [3, 4, 39–44]. All these rules share the condition that

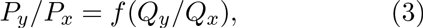

where *P*_*x*_, *P*_*y*_ are the probabilities to choose *x* and *y* respectively, *f* is any monotonically increasing function and *Q*_*x*_ ≥ 0, *Q*_*y*_ ≥ 0 are the estimated qualities for options *x* and *y* respectively. When more than two options exist, a relative decision rule must fulfil Equation 3 for every pair of options.

Relative decision rules give superaggregation in adversity whenever Equation 2 is fulfilled. If there are only two options (*x* and *y*), we must have *P*_*x*_ = 1 − *P*_*y*_. Therefore, decreasing *P*_*y*_ necessarily means decreasing *P*_*y*_/*P*_*x*_, so

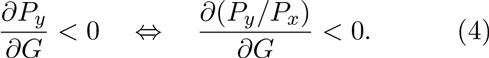

And from Equation 3 we have that

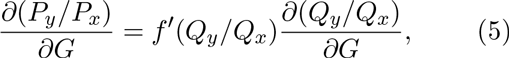

where *f*′(·) is the derivative of *f* with respect to its argument, which is always positive because *f* is by definition a monotonically increasing function. Therefore,

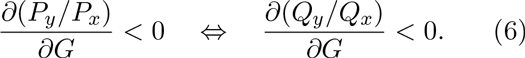
Putting together equations 4 and 6, superaggregation in adversity will take place whenever

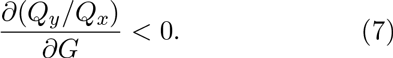

If the choice is among more than two options,asimilar condition is suficient (but not necessary) to produce superaggregation in adversity. The probability to choose the majority option must decrease if the ratio of this probability with respect to all other probabilities decreases:

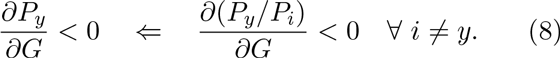

With this and Equation 6 (which is valid for any two options), we have that

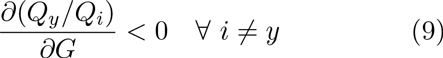

is a suficient condition for superaggregation in adversity when there are more than 2 options.

### Absolute decision rules

We say that a decision rule is absolute when the probability to choose option *y* is of the form

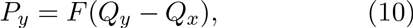

where *Q*_*x*_, *Q*_*y*_ are the qualities of options *x* and *y*, and *F* is any monotonically increasing function.

Absolute decision rules give superaggregation in adversity whenever Equation 2 is fulfilled. From Equation 10,

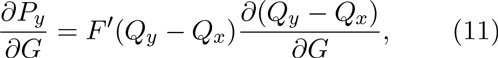

where *F* ′(·) is the derivative of *F* with respect to its argument, which is always positive because *F* is monotonically increasing. Therefore, 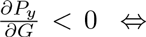 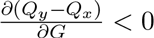 so superaggregation in adversity will take place whenever

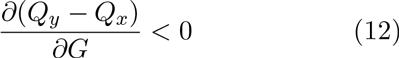

Absolute decision rules rule may emerge for example if the subject chooses *y* when *Q*_*y*_ > *Q*_*x*_ + *η*, with *η* a random number: Let *ρ*(*η*) be the probability density function of *η*, and *F*(*η*) its cumulative distribution. Then, the probability to choose *y* is equal to the probability to draw a value of *η* lower than *Q*_*y*_ − *Q*_*x*_,

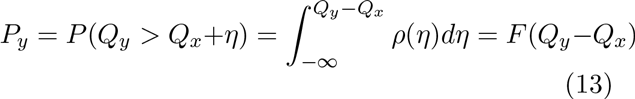

*F* is a cumulative distribution, which is always monotonically increasing. In Figures 1 and 2 we have used

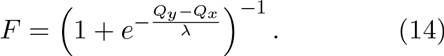

This function is the cumulative distribution of the logistic distribution, whose probability density function is 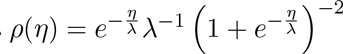 We have chosen this distribution because its cumulative density function has a simple analytical form; all results hold regardless of the probability distribution that we choose.

### Bayesian estimation model

This model is equivalent to our previously published model [4], with some changes in notation. We consider an individual choosing between two options, *x* and *y*. Each option may be good or bad, and the deciding individual estimates the probability that each option is good using both its private information and the behaviors of other individuals. For option *x*, we represent this probability as *P*(*X*|*B*,*C*) where *X* stands for ‘*x* is good’, *B* represents the behaviors of other individuals and *C* represents the private information of the deciding individual. Using Bayes’ theorem, we get

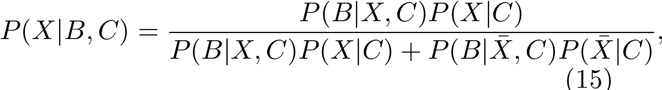

where 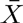 means ‘x is not good’ and 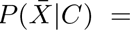 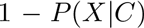. We define *G*_*x*_ = *P*(*X* |*C*), which is the privately estimated probability that *x* is a good option. Dividing numerator and denominator over the numerator, we get

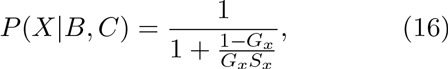

where 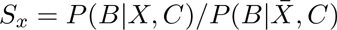 contains the social information. Our previous work develops the social term further adding some extra hypotheses in order to obtain an expression that depends on the number of individuals choosing each option [4]. This is not necessary here because our results are independent on the exact form of the social term; we only need to assume that *S_x_* increases when the behaviors of the other individuals indicate that *x* is good.

The quality of each option is given by the probability that it is good *Q*_*x*_ = *P*(*X*|*B*; *C*), and similarly for option *y*. In the case of symmetric private information, *G* ≡ *G*_*x*_ = *G*_*y*_, we have

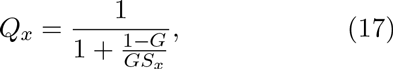

and similarly for option *y*.

When combined with a relative decision rule, the Bayesian model will show superaggregation in adversity when 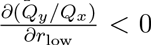 (Equation 1). From Equation 17,

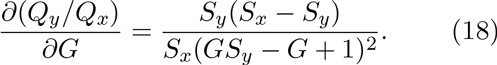

This expression is always negative because squared terms are always positive, by definition *S*_*x*_ > 0 and *S*_*y*_ > 0, and *S*_*y*_ > *S*_*x*_ because *y* is the majority option. Therefore, the Bayesian estimation model with a relative decision rule always produces superaggregation in adversity.

When combined with an absolute decision rule, the Bayesian model will show superaggregation in adversity when 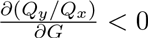 (Equation 12). From Equation 17,

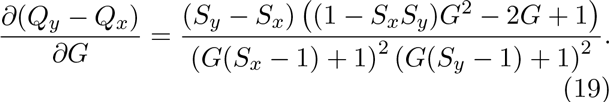

The denominator of this expression is always positive, and *S*_*y*_ − *S*_*x*_ is positive when *y* is the majority option. Therefore, the whole expression is negative when

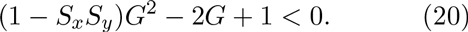

This inequality has only one solution for *G* > 0, which is

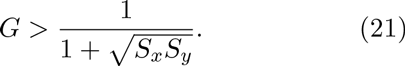

Thus, there is superaggregation in adversity in the range of high *G*, and the opposite effect in the range of low *G*. The higher *S*_*x*_*S*_*y*_, the wider the range for superaggregation in adversity.

This result matches the intuition that the upper bound is more important than the lower one because social information tends to move the estimation towards the upper bound. In principle an individual choosing option *x* can indicate both that *x* is a good option and/or that *y* is a bad one. Therefore, this choice can increase *S*_*x*_ in the same degree as it decreases *S*_*y*_. However, experimental data shows that social information usually has a positive net effect, meaning that an individual that chooses *x* increases *S*_*x*_ more than it decreases *S*_*y*_ (in our previous work [4] we defined parameter *k* to measure this effect. All experimental data, from three different species, were consistent with *k* < 1, meaning that social information has a positive net effect). When social information has a positive net effect, the product *S*_*x*_*S*_*y*_ increases as more individuals make choices. Therefore for large groups (or when behaviours are very informative) the range of superaggregation in adversity is very wide. For example, in our previous study [4] we found that for zebrafish *S*_*x*_*S*_*y*_ = 5^*n*^, where *n* is the total number of individuals that have already chosen one of the two options. Therefore, Equation 21 tells us that a group of 10 zebrafish will show superaggregation in adversity for *G* > 3 · 10^−4^, and a group of 15 for *G*> 6 · 10^−6^. As a reference, in our experiments (that corresponded to an intermediate level of adversity, with the fish in an unknown environment but without any direct threat) we found *G* ≈ 0.08 [4].

### Model with payoffs

Let *r*_low_ ≥0 be the reward provided by a bad option, and *r*_high_ ≥ 0 the reward provided by a good one, with *r*_low_ ≤ *r*_high_. The estimated quality of option *x* is its expected payoff

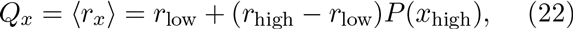

and similarly for option *y*. *P*(*x*_high_) is the estimated probability that *x* has a high payoff (*r*_high_), and Equation 22 takes into account that *P* (*x*_low_) = 1 − *P*(*x*_high_). The private estimate of the quality of the options is

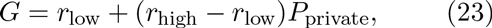

where *P*_private_ is the privately estimated probability that an option contains a high reward (equal for all options, by hypothesis). A change in private information may translate in three different changes in the parameters: (1) a change in the probability (*P*_private_), (2) a change in the lower payoff (*r*_low_) and
(3) a change in the higher payoff (*r*_high_). The first type of change will depend on the specific model we use to estimate the probabilities (we may use a Bayesian model as in previous sections, or any other model). Here we will consider the other two cases, in which the new private information changes the values of the rewards that the subject expects to obtain.

First let us consider the case in which adversity means a decrease in the lower reward (*r*_low_), and the case of a relative decision rule (Equation 3). From Equation 23 we have that 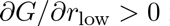 in all cases. Therefore, any derivative with respect to *G* will have the same sign as a derivative with respect to *r*_low_, and Equation 1 is now equivalent to 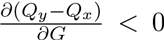. We compute this derivative from Equation 22, getting

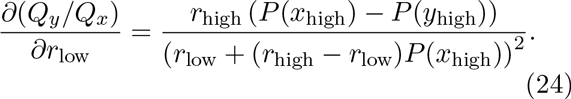

The denominator is always positive because it is squared, and *r*_high_ is positive by definition. *P*(*y*_high_) > *P*(*x*_high_) because *y* is the majority option, so the derivative is always negative; superaggregation in adversity always takes place.

We find the same result for an absolute decision rule. To test for superaggregation in adversity we use Equation 12, deriving with respect to *r*_low_ instead of *G* (as explained in the previous paragraph). We get

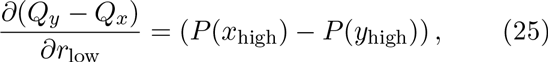

which is always negative because *P*(*y*_high_) > *P*(*x*_high_) when *y* is the majority option. Therefore, cohesion always increases with adversity.

In contrast, cohesion decreases in adversity when the higher payoff changes, keeping all other parameters constant. This has a simple intuitive explanation: in adversity *r*_high_ goes down, becoming more similar to *r*_low_. Therefore, the difierence between choosing correctly and choosing incorrectly decreases, making any decision weaker. Mathematically, for relative decision rules we evaluate the condition in Equation 1 (in this case deriving with respect to rhigh instead of *G*), finding

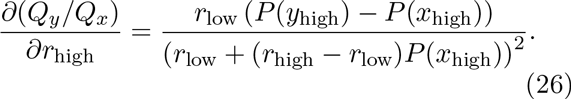

The denominator of this expression is always positive, as is rlow by definition. And *P*(*y*_high_) > *P*(*x*_high_) because *y* is the majority option, so the derivative is always positive, meaning that cohesion decreases in adversity when using a relative decision rule. Now we consider an absolute decision rule (Equation 12):

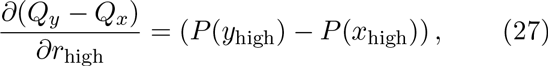

which is always positive because *P*_*y*_ > *P*_*x*_ when *y* is the majority option. Therefore, cohesion decreases in adversity.

### Model for the cards experiment

The experimental design presented social information as a separate option (Figure 4b), so we have implemented the simplest model that mimics this condition: the subject assumes that the other person (the professor or the student) has a fixed probability of making a correct choice (ξ). And the probability of making a correct choice by choosing a card is *n*_red_/8, where *n*_red_ is the number of red cards. We then consider a 9-choice scenario: the 8 cards plus the social option. As decision rule we use probability matching, where the probability to choose one option is proportional to its corresponding probability of success. Then, the probability to rely on the other’s opinion is

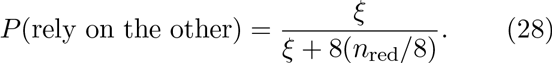

We fit the parameter ξ to the experimental data, getting ξ = 0:624 for the professor and ξ = 0:328 for the student.

## Acknowledgements

We are very grateful to Sara Arganda, Barrett Deris, Francisco Ferrero-Romero and Raphael Jeanson for fruitful discussions, and to Trevor I. Case for clarifications about the data from ref. [47].

## SUPPLEMENTARY MATERIAL

### 1 Alternative Bayesian model

The Bayesian model used in the main text assumes that the quality of each option is estimated independently of the other. Here we remove this approximation, arriving to a different and more complex model. Regarding superaggregation in adversity, we find the same conclusions as for the simpler model presented in the main text.

#### 1.1 Derivation of the estimated qualities

Let us consider a choice between two options, *x* and *y*. Each option can be good or bad; we will write *X* to denote ‘*x* is good’ and 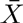 to denote ‘*x* is bad’, and similarly for option *y*. The pair of options can be in four possible states: both options are good (*XY*), both options are bad 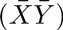, or one option is good and the other is bad 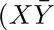 and 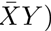. We calculate the probability for each of these states, using both private information (*C*) and the behaviours of the other individuals (*B*). For example, the probability that option *x* is good and option *y* is bad, using Bayes’, theorem, is

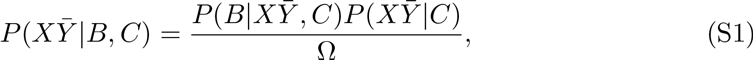

where

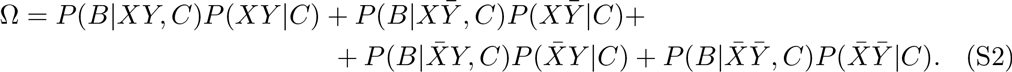

The term 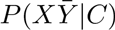 contains the private information about the state of both options, and the term 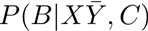 contains the social information. If we assume that the two options can be good or bad independently, we have 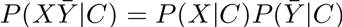. Now we define *G*_*x*_ = *P*(*X*|*C* and *G*_*y*_ = *P*(*y*|*C*, so 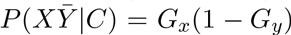. If we further assume that private information is symmetrical (*G*_*x*_ = *G*_*y*_ = *G*),
Equation S1 becomes 
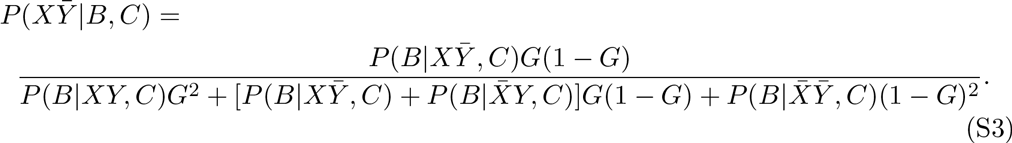

The four probabilities *P*(*B*|*XY*, *C*), 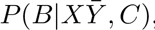, 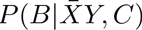 and 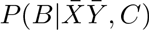 parametrize the available social information. Because they must sum one, we only have three free parameters. It is therefore more useful to define 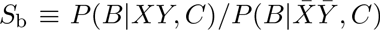, 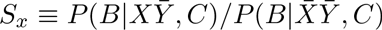 and 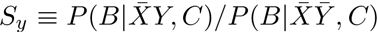, and write Equation S3 as

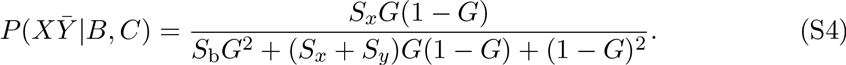

The probabilities for the other three states can be derived in the same way:

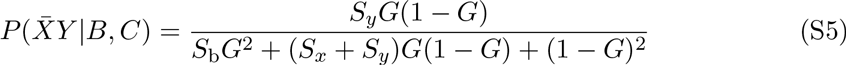

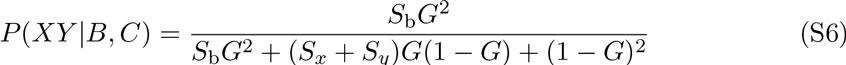

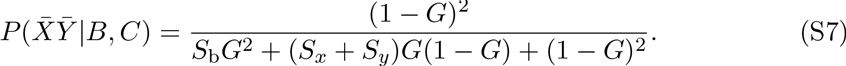

Now we define the quality of *x* as the probability that *x* is good (and the same for option *y*), getting

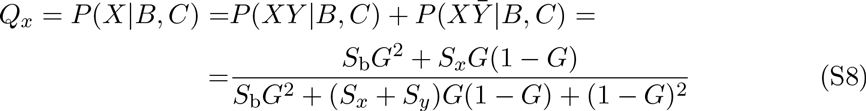

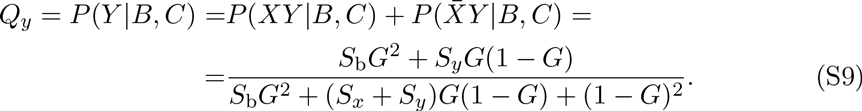

#### 1.2 Effect of a relative decision rule

We assume that y is the majority option. If the decision rule is relative (Equation 3 of the main text), superaggregation in adversity will take place when 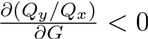 (Equation 7 of the main text). From Equations S8 and S9, 
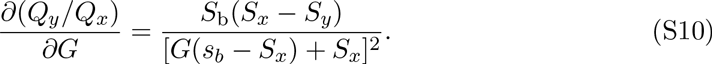

The denominator of this expression is always positive because it is squared. *S*_*b*_ is always positive because it is a ratio of probabilities. And *S*_*y*_ > *S*_*x*_ because *y* is the majority option, so this derivative is always negative. Therefore, there is superaggregation in adversity for any values of the parameters.

#### 1.3 Effect of an absolute decision rule

If the decision rule is absolute (Equation 10 in the main text), superaggregation in adversity will take place when 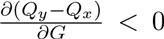 (Equation 12 of the main text). From Equations S8 and S9,

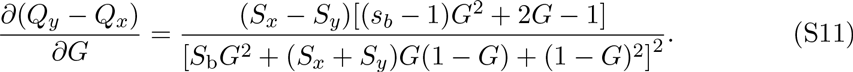

The denominator is always positive because it is squared. *S*_*x*_ − *S*_*y*_ is always negative because *S*_*y*_ > *S*_*x*_ when *y* is the majority option. Therefore, the sign depends on the sign of (*s*_*b*_ − 1)*G*^2^ + 2*G* − 1. This polynomial has a single root between 0 and 1 at 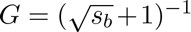. Therefore, the derivative is negative when 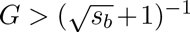, recovering the same result as for the Bayesian model in the main text: there is superaggregation in adversity in the regime of high *G*, and the opposite effect in the regime of low *G*.

### 2 The selfish herd hypothesis in the quality landscape

We assume that the available space is divided in *M* possible locations. The *i*-th location is occupied by *n*_i_ individuals (*i* = 1 … *M*). The quantities n_i_ do not count the focal individual, which starts from any given location. A predator may arrive to any location with probability 1 − *G* (we define it in this way to keep the convention that *G* decreases when conditions become adverse). If the predator arrives, it will eat one of the individuals in that location, chosen at random. We define the quality of each option as the probability that the focal individual survives after choosing that location, so for location *k* we have

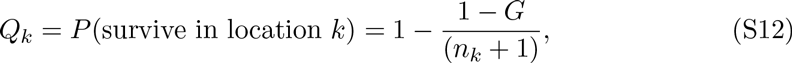

where *n*_*k*_ + 1 is the number of individuals in option *k*, assuming that the focal individual chooses it and that no other individual moves in the current round.

Figure S1a shows the trajectory of this estimation rule for the case of two locations (*M* = 2), when private information modifies the value of *G*. The probability of following the majority increases in adversity both for the relative decision rule (Figure S1b, solid line) and for the absolute one (Figure S1b, dashed line).

### 3 Spatial model

In order to simulate animals in motion, we discretize the space in pixels (for 2D simulations) or voxels (for 3D simulations). In each iteration, each individual chooses one pixel and accelerates towards it, up to a maximum acceleration (*a*_max_) and never exceeding a maximum velocity (*v*_max_). The probability to choose a given pixel is proportional to the probability that it is a good place according to a model based on Bayesian estimation [1]

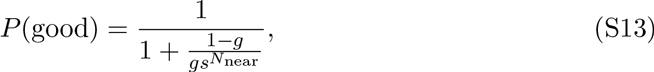

where *g* is the probability that each location is good from non-social information only. Here we define the qualities as *Q* = *P*(good), so *g* is also the privately estimated quality (*G* = *g*). *N*_near_ is the number of individuals within a certain radius *r*_influence_ from the pixel, and s is a parameter measuring the reliability of the other individuals. We also incorporate a limited field of view for each individual, by assuming that an individual can only choose pixels within a certain radius from itself, *r*_view_ (this parameter is needed to make the model computationally tractable even when the individuals are not restricted to a finite region).

**Figure S1:**
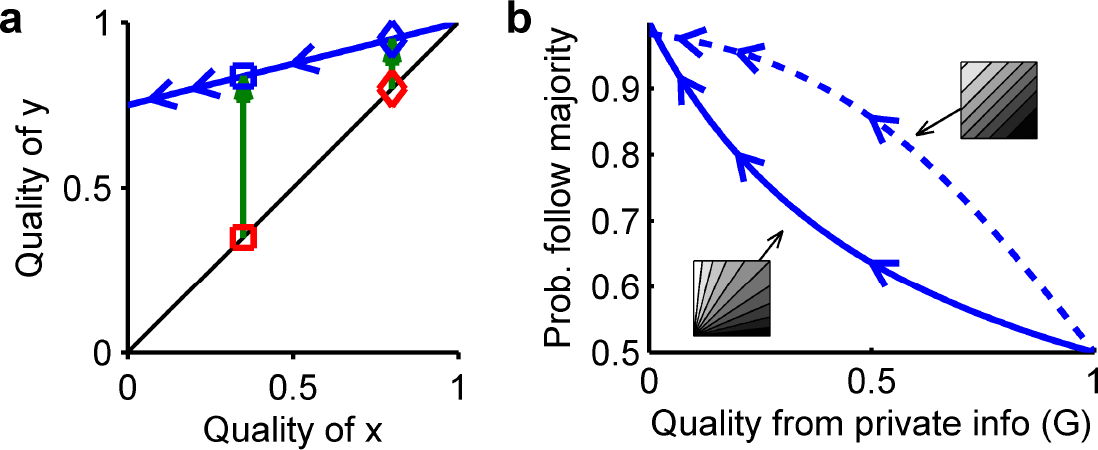
Selfish herd in the quality landscape. **a.** Change in estimated qualities for a selfish herd model (Equation S12). Red: private estimate for favorable conditions (diamond) and adverse conditions (square). Green arrows: contribution of the social information. Blue: trajectory of the final estimated qualities. **b.** For the decision model in (a), probability of choosing the majority option (*y*), as a function of the privately estimated quality (*G*, higher values indicate more favourable conditions). Solid line: relative decision rule. Dashed line: absolute decision rule.

For the results in Figure 3a,b and Movie S1, we chose the parameters *a*_max_ = 0.1, *v*_max_ = 0.5, *r*_influence_ = 5, *r*_view_ = 100, *s* = 3, and G as indicated in Figure 3b (space units are in pixels, and time units in iterations).

For the case of predator avoidance the model confirmed our prediction of increased cohesion in adversity, but was not realistic: in many conditions, when the predator appears individuals cluster together but fail to run away from it (the social attraction overcomes the repulsion from the predator). In order to obtain more realistic results, we added another factor: an individual indicates that the location towards which it is heading (rather than its current location) is a good place. To account for this, we centre the circle of influence of each individual at the position where it will be in *t*_prediction_ steps in the future (assuming it will keep constant direction and speed), rather than at its current location. With this addition, animals not only cluster together when the predator appears, but also tend to align with each other along the optimal scape route (Figure 3c,d, and Movie S2; simulations with the same parameters as in previous paragraph, but with *t*_prediction_ = 15).

In fact, alignment arises in adversity when individuals pay attention to future positions rather than current ones (*t*_prediction_ > 0), even if there is no predator present (Movie S3; simulations with same parameters as for Movie S1, but with *t*_prediction_ = 15). Thus, in our model the group does not need the asymmetry created by the predator to reach a consensus direction. But if there is a predator, the consensus direction will head away from it.

Figure 3e shows the average distance between individuals after a simulation with a high value of *G* (favorable conditions, *y* axis) vs. the average distance after a simulation with a low value of *G* (adverse conditions, *x* axis). Each average distance comes from 50 simulations with identical parameters and random initial positions. Each simulation lasts 200 iterations. For each combination of parameters, we run simulations with all values of *G* ∈ {10^−3^, 10^−2.5^,10^−2^,…, 10^0^}. Then we plot the results of all pairs of *G* (always with the smaller *G* in the *x* axis, and the greater *G* in the *y* axis). To generate combinations of parameters, we drew random numbers uniformly distributed in the following intervals: *a*_max_∈ [0.1, 5], *v*_max_ ∈ [1, 50], *r*_influence_ ∈ [5,30], *r*_view_ ∈ [10, 50], *t*_prediction_ ∈ [0, 5], *s* ∈ [2,20]. For each combination of parameters, we run the simulation both in 2D and 3D.

To generate Figure 3f we followed the same procedure as for Figure 3e, but instead of a Bayesian decision rule (Equation S13) we used an additive rule where the quality of a given pixel is

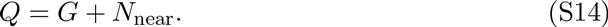

*G* is again the privately estimated quality (but note that it is not a probability any more), and *N*_near_ is the number of individuals near the pixel (as above). We drew random parameters from the same intervals mentioned in the previous paragraph, except for *s* (this parameter does not exist in this model). Also, in this case *G* ∈ {0,0.1,0.5,1, 2, 5,10}. All other details are as described in the previous paragraph.

To generate Figure 3g we followed the same procedure as for Figure 3e, but the quality of each pixel is given by a model with explicit payoffs (Equation 22 in main text). To compute the probability that each pixel has a high payoff we use the Bayesian model above (so *P*(*x*_high_) in Eq. 22 is equal to *P*(good) in Eq. S13). Adverse conditions now mean a low value of *r*_low_, so we keep *g* = 0.05 and *r*_high_ = 1 fixed, and we change *r*_low_ ∈ {0, 0.001, 0.01, 0.1, 0.2, 0.4, 0.8}.

To reproduce the experimental data in Figure 4a, we adjusted the model parameters to mimic the experimental conditions [2]. The experiments were performed in a 100x100x10 cm tank, and the experimenters considered that two fish belonged to the same group when they were closer than 16 cm. Assuming a scale of 1 pixel/cm, we simulated 10 individuals in a closed 2D space of 100x100 pixels, and set *r*_influence_ = 16 pixels. When performing the analysis, we consdered that two fish belonged to the same group when they were within 16 pixels. Then wekov searched parameter space manually and non-systematically, finding a good fit for *a*_max_ = 1, *v*_max_ = 12, *r*_view_ = 30, *s* = 10, *t*_prediction_ = 0, and *G* as in Figure 4a.

